## Supplementary Materials for "Mapping the architecture of the initiating phosphoglycosyl transferase from *S. enterica* O-antigen biosynthesis in a liponanoparticle"

[b]Thermo Fisher Scientific, San Jose CA 95134, USA.

**ORCID IDs:**

BI: 0000-0002-5749-7869

RV: 0000-0003-0550-5545

WJ: 0000-0003-2322-0500

YH: 0000-0002-5565-5579

AJA: 0000-0003-4871-4283

GJD: 0000-0002-6555-8350

Table S1 Screening chemical crosslinkers for the detection of S. enterica WbaP oligomers. S. enterica WbaP in SMALP was reacted with a panel of lysine-reactive crosslinkers with variable chemistries, solubilities, and lengths. Samples were analyzed by Western blot to detect the presence of crosslinked oligomers. Crosslinking efficiency was annotated as high (>30 % total protein crosslinked to dimer), moderate (<30 % total protein crosslinked to dimer), or none (no protein crosslinked to dimer).

| Crosslinker | Chemistry | Membrane permeable | Water soluble | Spacer Length (Å) | *Se* WbaP crosslinking efficiency |
|---|---|---|---|---|---|
| DMA | imidoester | yes | yes | 8.6 | none |
| DMS | imidoester | yes | yes | 11 | none |
| BSOCOES | NHS ester | yes | no | 13 | moderate |
| DSP | NHS ester | yes | no | 12 | high |
| DTSSP | NHS ester | no | yes | 12 | high |

Table S1 **Cryo-EM data collection, refinement and validation statistics**

| | *S. enterica* WbaP in SMALP #1 (EMD-41042) (PDB 8T53) |
|---|---|
| **Data collection and processing** | |
| Microscope / Detector | Krios G3i, K3 |
| Magnification | 105,000X |
| Voltage (kV) | 300 |
| Electron exposure (e–/Å$^2$) | 50.13 |
| Defocus range (μm) | 0.75 - 2.5 |
| Pixel size (Å) | 0.87 |
| Total movies (no.) | 4,841 |
| Symmetry imposed | C2 |
| Initial particle images (no.) | 1,266,538 |
| Final particle images (no.) | 196,663 |
| Map resolution (Å) | 4.28 |
| FSC threshold | 0.143 |
| Map resolution range (Å) | |
| **Refinement** | |
| Initial model used | *Se* WbaP AlphaFold dimer model |
| Model composition | |
| Non-hydrogen atoms | 6660 |
| Protein residues | 796 |
| Ligands | 0 |
| *B* factors (Å$^2$) | |
| Protein (min/max/mean) | 65.79/728.40/310.78 |
| Ligand | n/a |
| R.m.s. deviations | |
| Bond lengths (Å) | 0.004 |
| Bond angles (°) | 0.944 |
| Validation | |
| MolProbity score | 1.59 |
| Clashscore | 11.81 |
| Poor rotamers (%) | 0 |
| Ramachandran plot | |
| Favored (%) | 98.19 |
| Allowed (%) | 1.81 |
| Disallowed (%) | 0 |
| **Model vs Data** | |
| CC (mask) | 0.45 |
| CC (box) | 0.45 |
| CC (peaks) | 0.23 |
| CC (volume) | 0.45 |

Table S3 NanoDSF nucleotide ligand screen for soluble WbaP DUF truncation.

| Sample | Ligand | Ligand Concentration | TM °C | ΔTM |
|---|---|---|---|---|
| Blank | None | N/A | 48.35 ± 21 | |
| AcCoA | acetyl coenzyme A | 200 µM | 47.95 ± 0.17 | -0.4 |
| AMP | adenosine 5'-monophosphate | 200 µM | 48.10 ± 0.07 | -0.25 |
| ATP | adenosine 5'-triphosphate | 200 µM | 47.98 ± 0.08 | -0.37 |
| CMP | cytidine 5'-monophosphate | 200 µM | 48.11 ± 0.30 | -0.24 |
| CoA | coenzyme A | 200 µM | 48.02 ± 0.05 | -0.33 |
| CTP | cytidine 5'-triphosphate | 200 µM | 48.14 ± 0.35 | -0.21 |
| dTTP | 2'-deoxythymidine 5'-triphosphate | 200 µM | 48.86 ± 0.21 | 0.51 |
| FAD | flavin adenine dinucleotide | 200 µM | 49.06 ± 0.18 | 0.71 |
| FMN | flavin mononucleotide | 200 µM | 48.35 ± 0.09 | 0 |
| GDP | guanosine 5'-diphosphate | 200 µM | 48.19 ± 0.04 | -0.16 |
| GTP | guanosine 5'-triphosphate | 200 µM | 48.00 ± 0.28 | -0.35 |
| NAD | nicotinamide adenine dinucleotide | 200 µM | 48.04 ± 0.01 | -0.31 |
| NADH | nicotinamide adenine dinucleotide | 200 µM | 48.54 ± 0.09 | 0.19 |
| NADP | nicotinamide adenine dinucleotide phosphate | 200 µM | 48.11 ± 0.04 | -0.24 |
| NADPH | nicotinamide adenine dinucleotide phosphate | 200 µM | 48.05 ± 0.26 | -0.3 |
| TMP | thymidine 5'-monophosphate | 200 µM | 48.40 ± 0.07 | 0.05 |
| UDP | uridine 5-diphosphate | 200 µM | 49.6 ± 0.11 | **1.25** |
| UDP-Gal | uridine 5′-(α-D-galactopyranosyl dihydrogen diphosphate) | 200 µM | 44.47 ± 0.12 | **-3.88** |
| UMP | uridine 5'-monophosphate | 200 µM | 48.44 ± 0.27 | 0.09 |
| UTP | uridine 5'-triphosphate | 200 µM | 48.41 ± 0.07 | 0.06 |

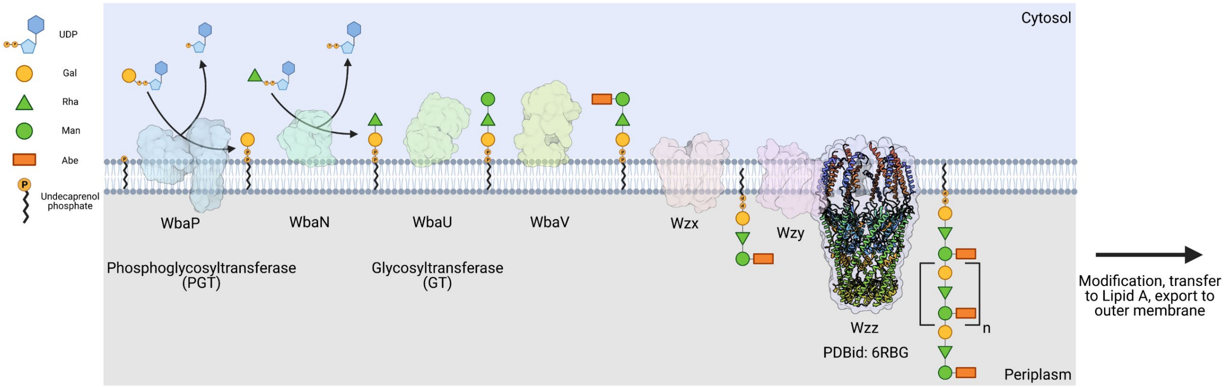


Figure S1 Biosynthesis of O-antigen repeat units in S. enterica servovar typhimurium. The pathway is initiated by the transfer of a phospho-Gal onto UndP, catalyzed by the LgPGT WbaP. A series of glycosyltransferases (GTs) add additional sugars onto the nascent RU. The repeat unit is then flipped across the membrane by a Wzx-class flippase, and RUs are polymerized through the coordinated action of the Wzy polymerase and Wzz chain-length regulatory protein.

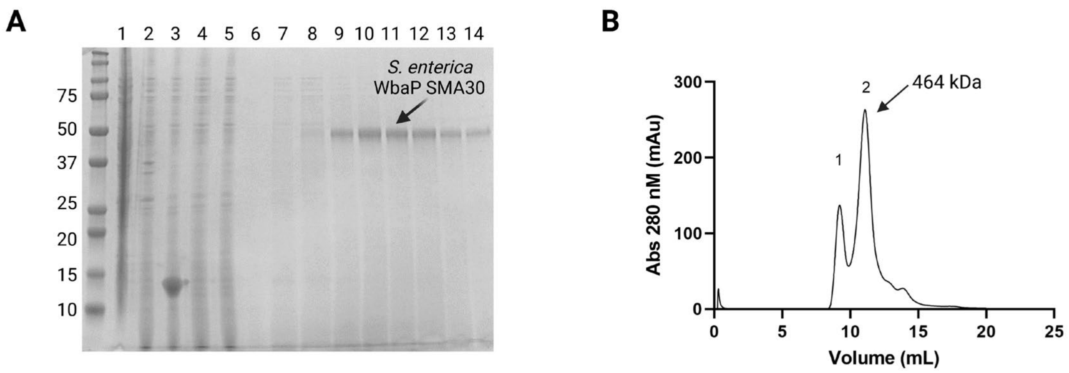


Figure S2 Purification and characterization of *S. enterica* WbaP in SMALP. **A**: Coomassie stained SDS-PAGE of large-scale WbaP purification. Lanes: 1: Lysate, 2: Cell envelope fraction (CEF), 3: SMA30-solubilized CEF, 4: StreptactinXT flowthrough, 5: Wash 1, 6: Wash 2, 7-14: Biotin elution. **B**: Purified WbaP separated on an Enrich SEC 650 column. Peak 1 corresponds to the void volume of the column, Peak 2 elutes at 11.09 mL, corresponding to a molecular weight of 464 kDa.

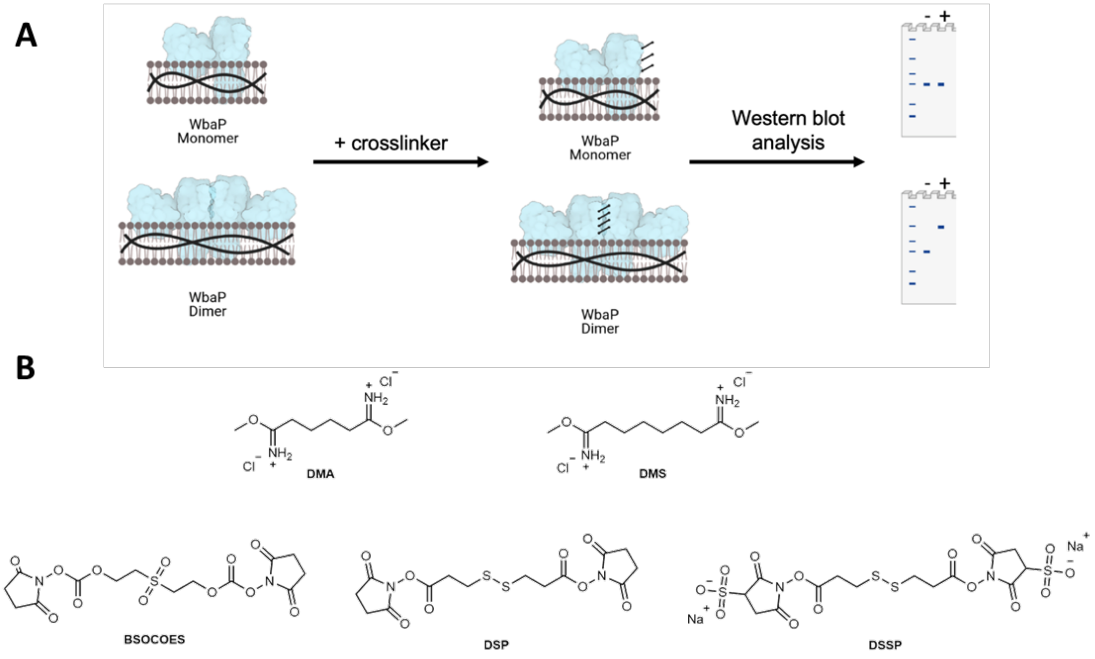


Figure S3 Crosslinking strategy for *S. enterica* WbaP in SMALP. **A**: A cartoon scheme depicting the results of crosslinking for either a WbaP monomer, or a WbaP dimer. Crosslinking efficiency is read out by SDS-PAGE or western blot. **B**: Panel of crosslinking compounds screened. DMA: dimethyl adipimidate, DMS: dimethyl suberimidate, BSOCOES: bis[2-(succinimidyloxycarbonyloxy)ethyl]sulfone, DSP: dithiobis(succinimidyl propionate), DTSSP: 3,3'-dithiobis(sulfosuccinimidyl propionate).

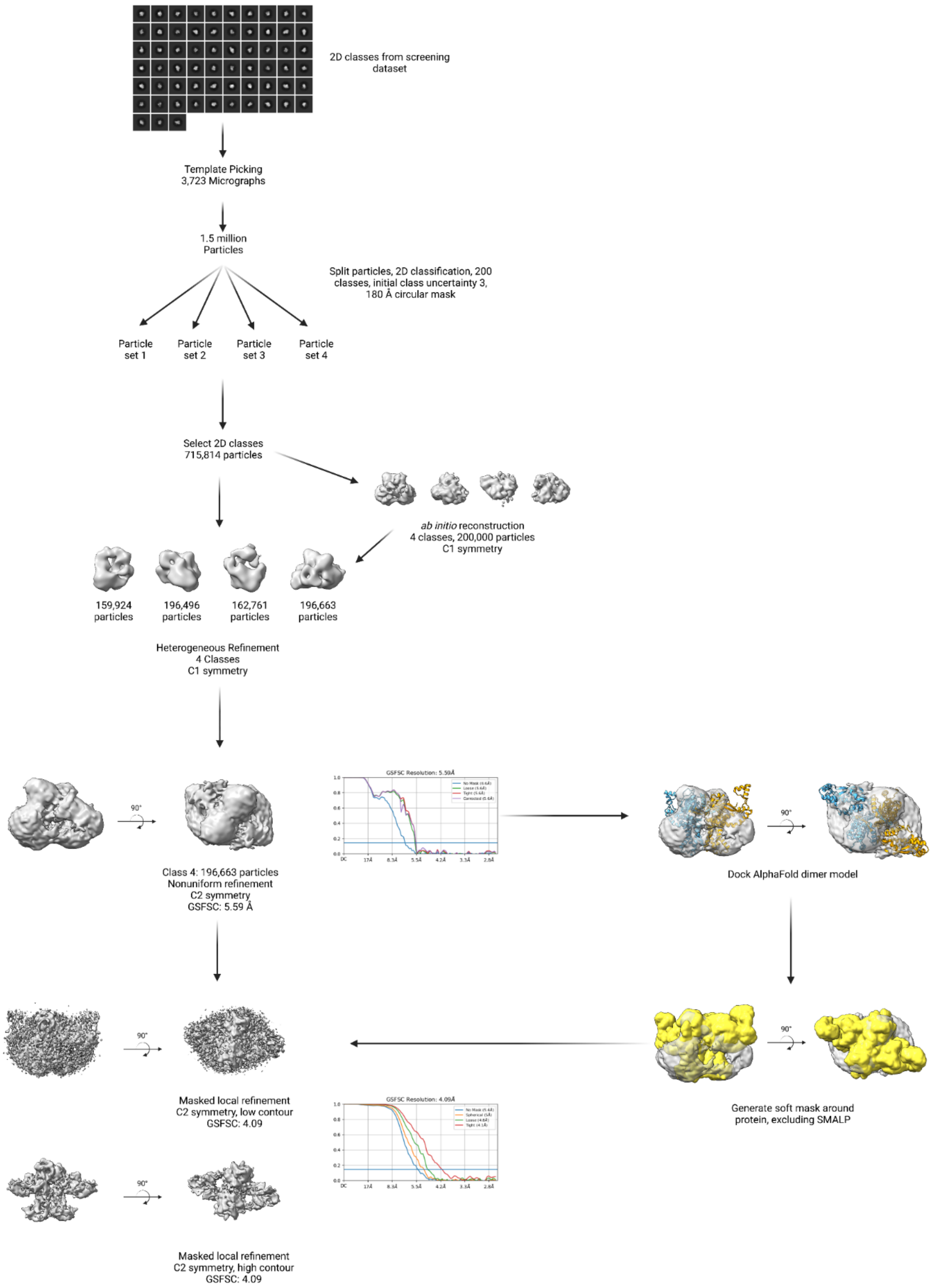


Figure S4 EM processing workflow

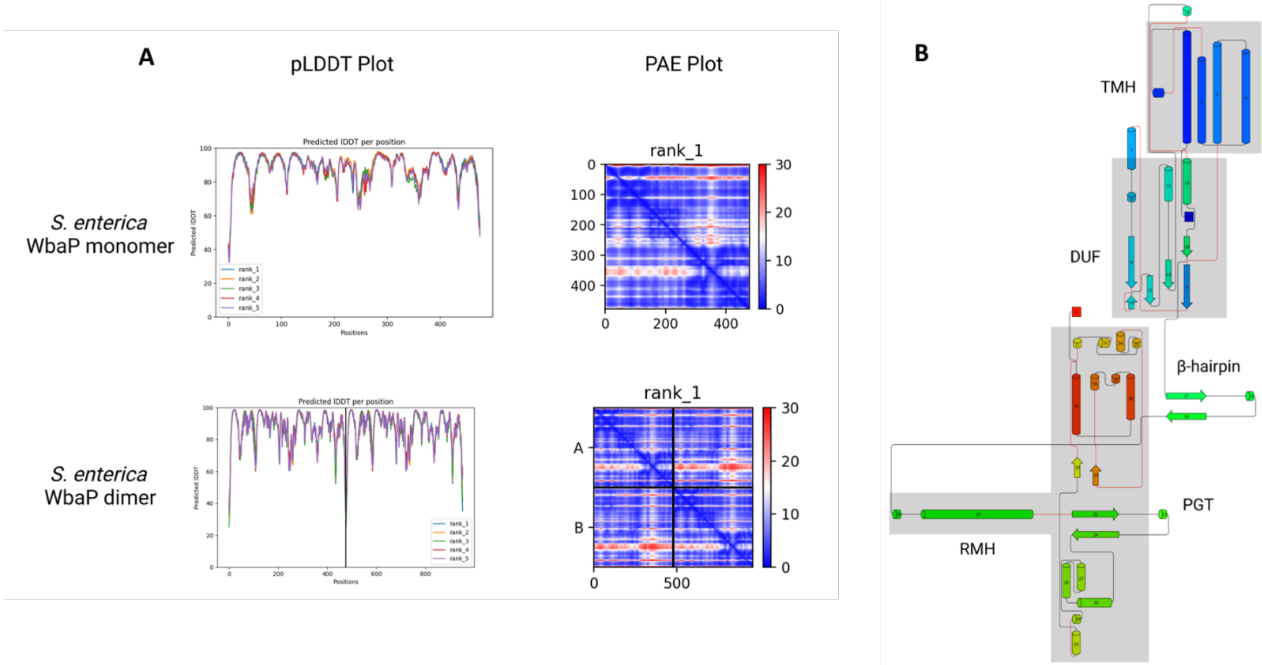


Figure S5 Analysis of *S. enterica* WbaP AlphaFold prediction. **A**: AlphaFold predicted local distance difference test (pLDDT) plots and predicted aligned error (PAE) plots for S. enterica WbaP monomer and dimer predictions. The overall confidence in both models is high. **B**: Topology diagram of AlphaFold S. enterica WbaP prediction. Structural domains shaded in gray. TMH: transmembrane helix, DUF: domain of unknown function, PGT: phosphoglycosyl transferase, RMH: re-entrant membrane helix

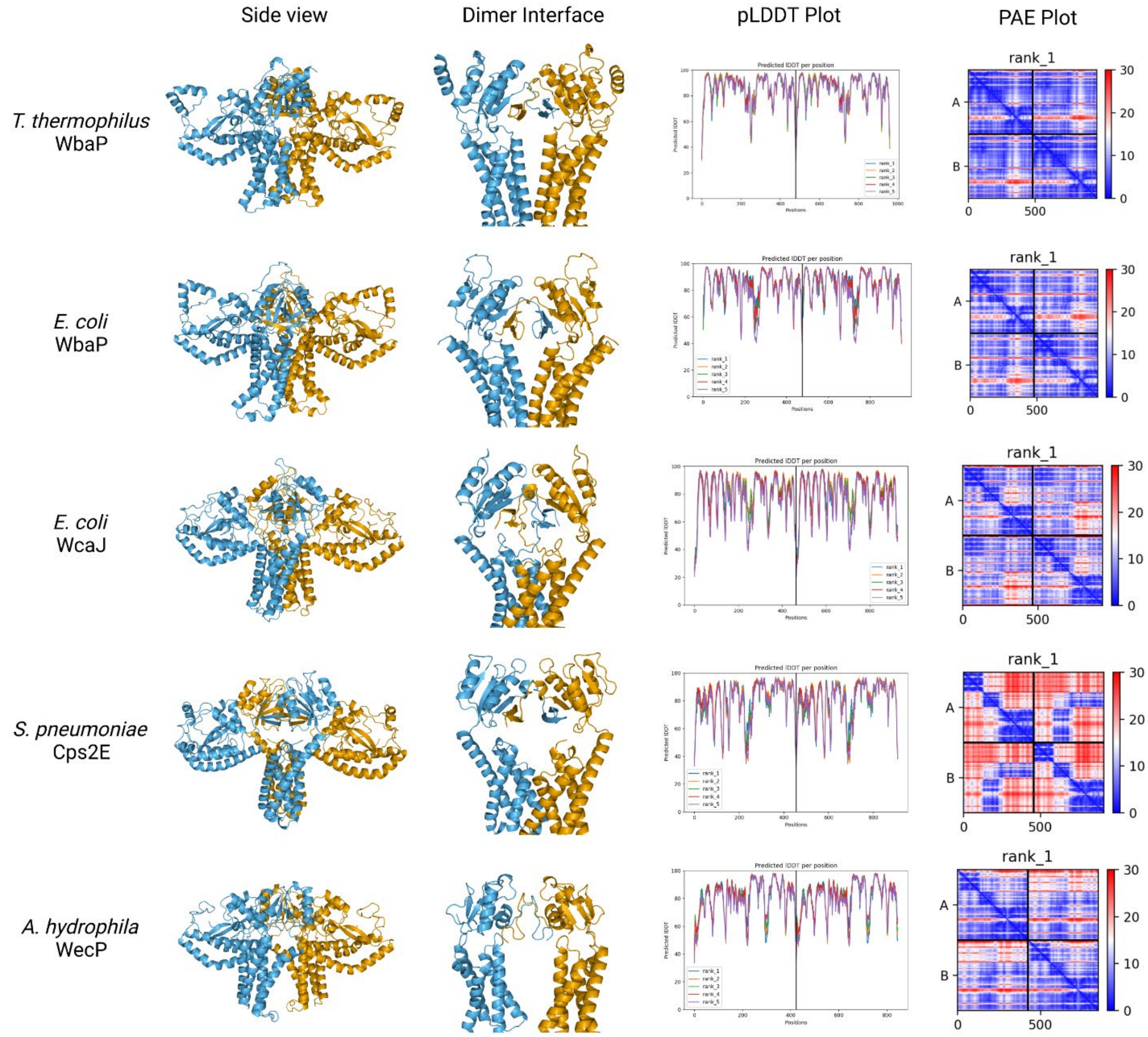


Figure S6 AlphaFold dimer predictions for various Lg-PGTs, along with predicted local distance difference test (pLDDT) plots and predicted aligned error (PAE) plots. β-hairpin-mediated domain swaps are observed for each prediction. While the estimated quality of predictions varies from protein to protein, the overall confidence in the models is satisfactory, with the majority of the regions of the proteins having a pLDDT score ≥ 70.

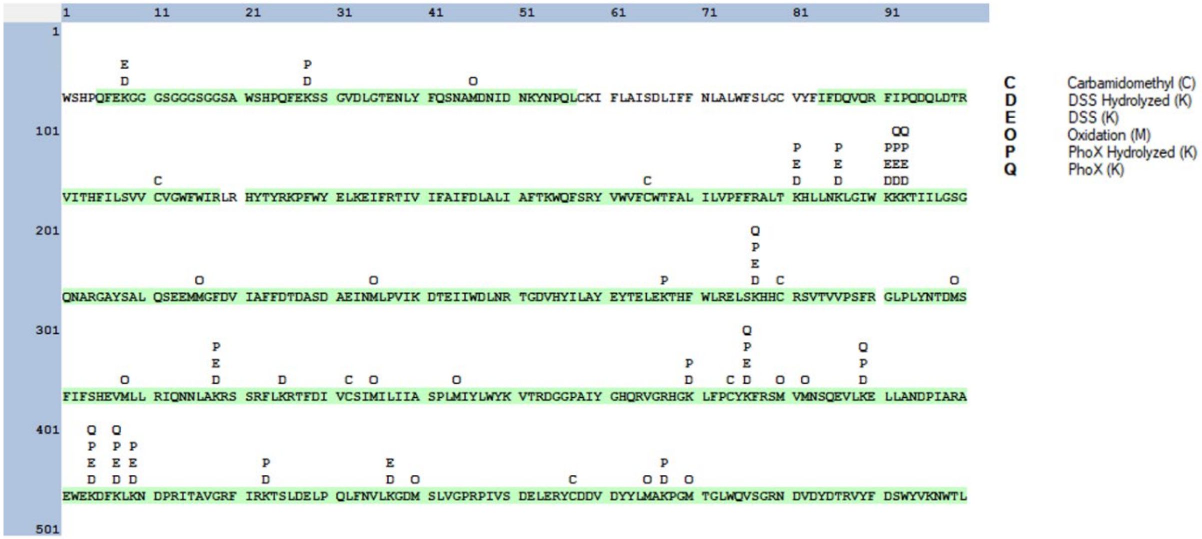


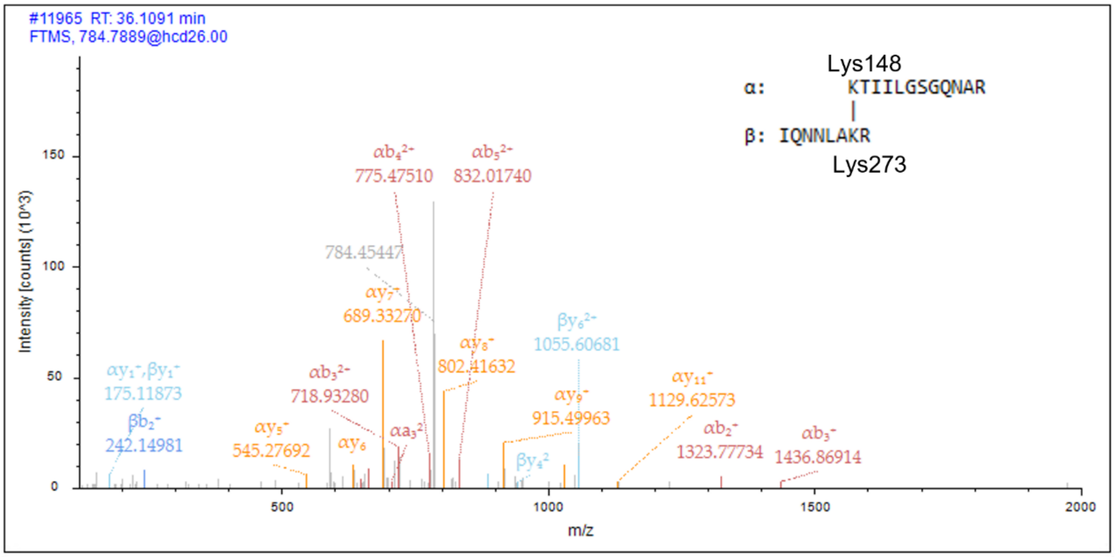


Figure S7 Crosslinking mass spectrometry analysis of *S. enterica* WbaP in SMALP. Right: Coverage and detected modifications across the WbaP construct primary sequence. Residues highlighted in green were detected in fragments after proteolytic degradation. Left: Example of MS/MS spectrum of the DSS inter-chain crosslinked fragment between Lys148 and Lys273.

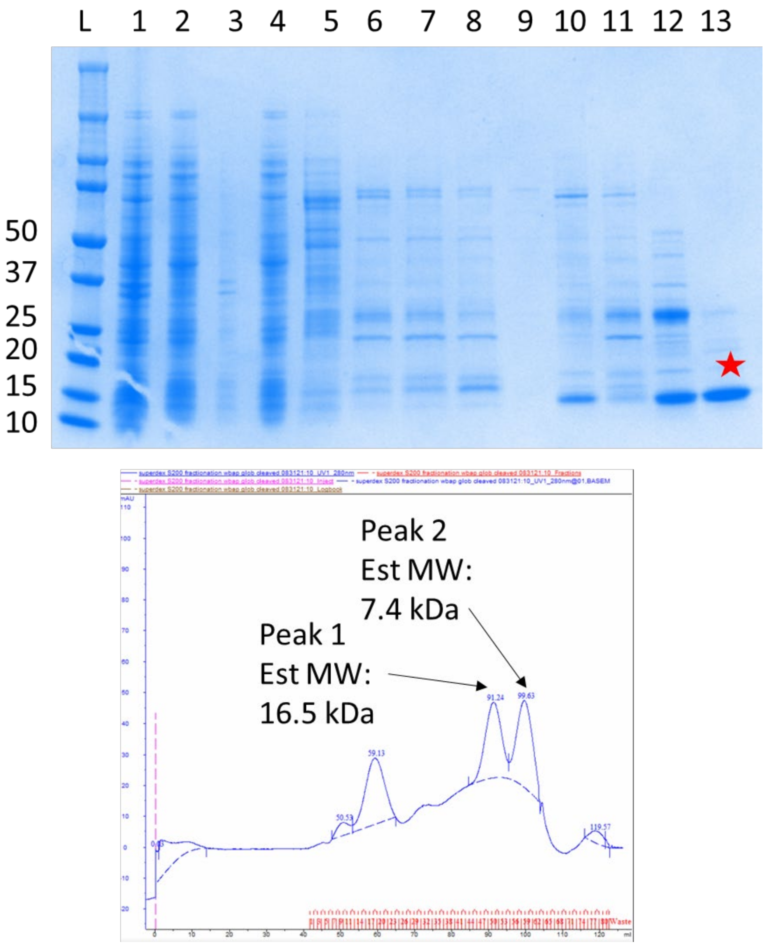


Figure S8 Purification of *S. enterica* WbaP soluble DUF truncation. **A**: SDS-PAGE gel monitoring purification. The final purified material indicated by red star. Predicted molecular weight: 14.83 kDa. Lanes: 1: Lysate, 2: Soluble, 3: Pellet, 4: flowthrough, 5: wash, 6-8: Elution, 9: Post-TEV flowthrough, 10: low imidazole Elution, 11: high imidazole elution, 12: S200 peak 1, 13: S200 peak 2. **B**. FPLC chromatogram of S200 separation.

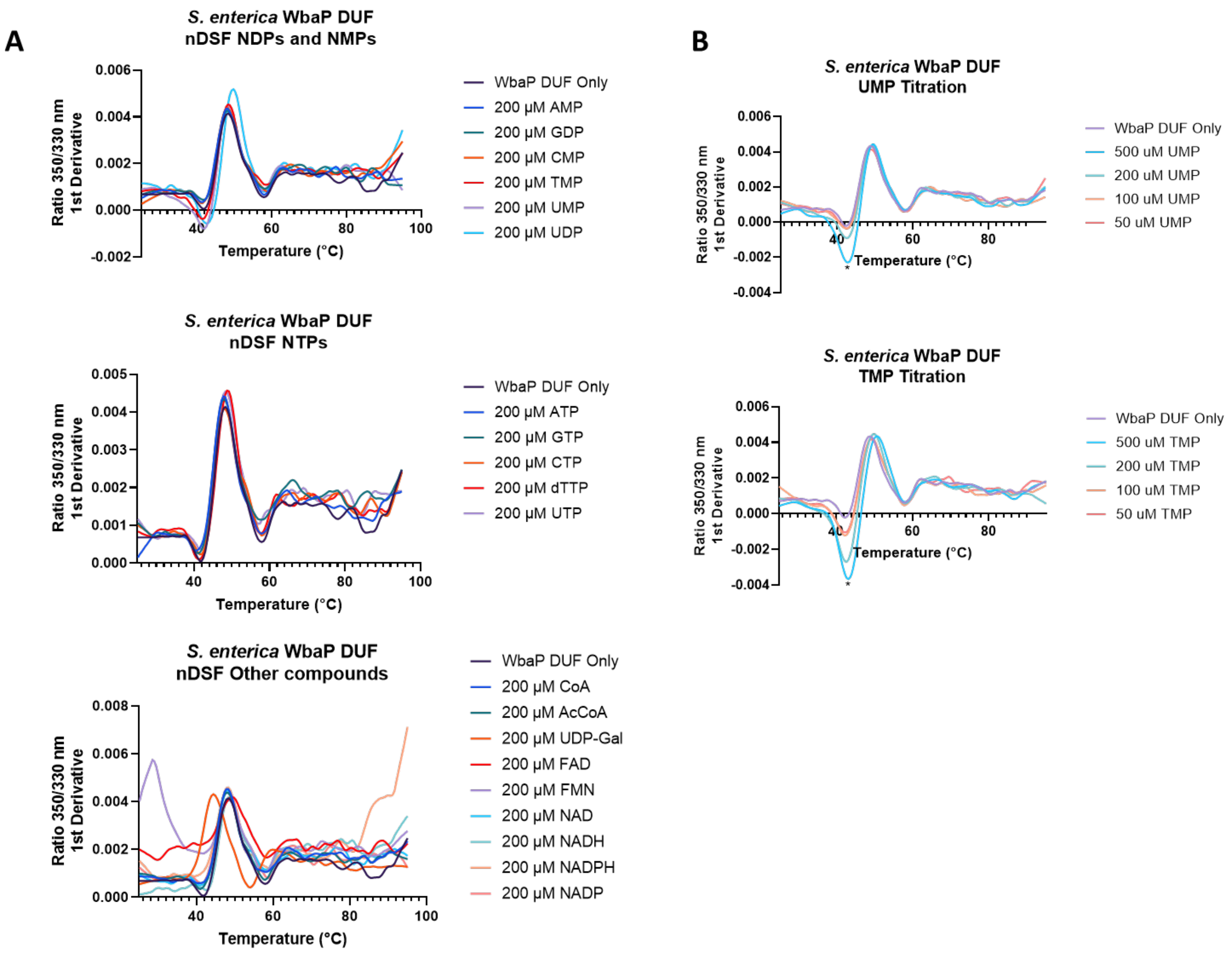


Figure S9 Nano differential scanning fluorimetry (nDSF) of purified soluble WbaP DUF truncation. **A**: Top: Incubation of DUF with selected nucleotide triphosphates (NTPs). Middle: Incubation of DUF with selected nucleotide diphosphates (NDPs) and nucleotide monophosphates (NMPs). Bottom: Incubation of DUF with additional putative small molecule ligands. **B:** Top: UMP titration, change in Trp/Tyr fluorescence 1$^{st}$ derivative indicated with asterisk. Bottom: TMP titration, change in Trp/Tyr fluorescence 1$^{st}$ derivative indicated with asterisk. See table S2 for tabulated shifts in melting temperature and full ligand names.

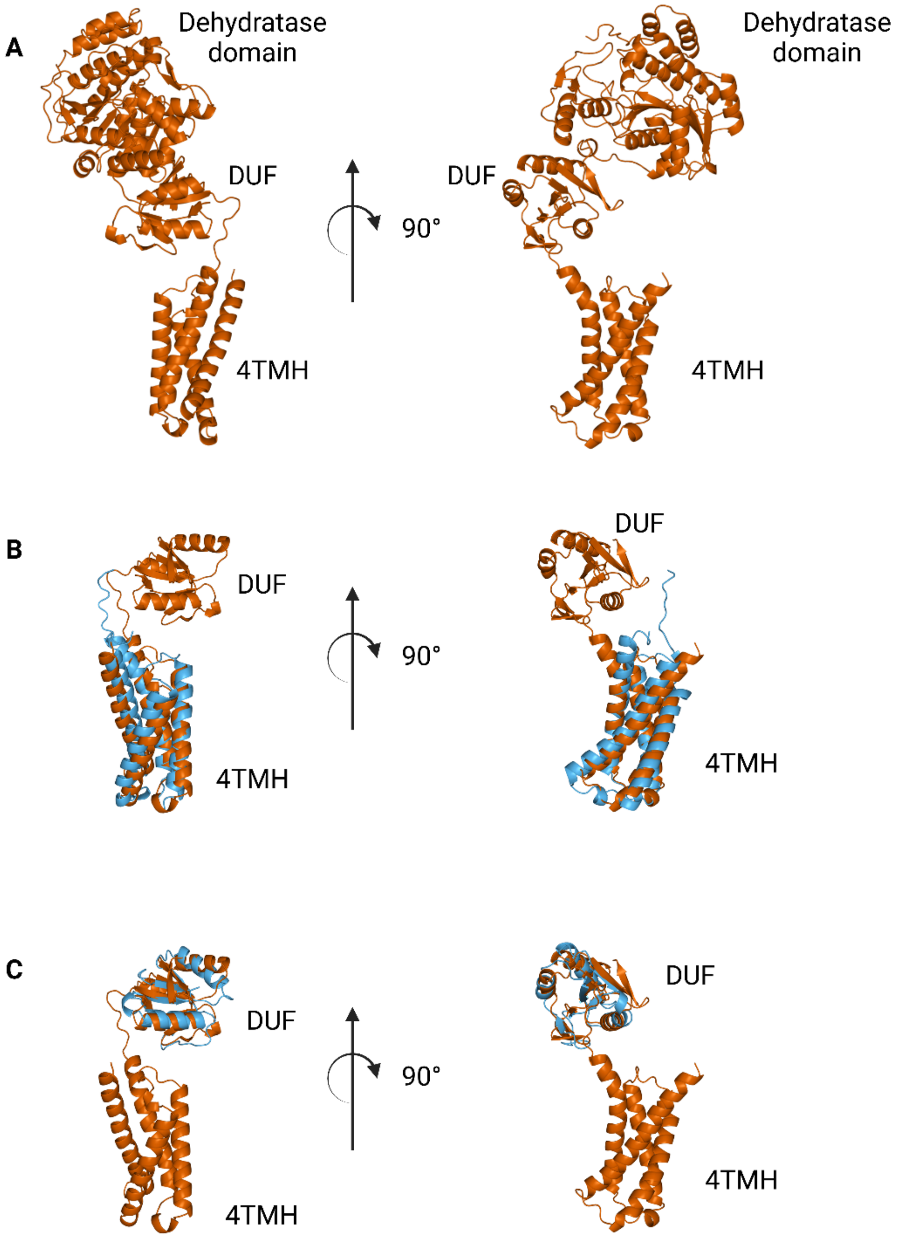


Figure S10 Comparison between N-terminal domains from *C. jejuni* PglF (Unitprot: Q0P9D4), a nucleotide sugar dehydratase, and S. enterica WbaP **A**: AlphaFold prediction of full-length PglF **B:** Superposition of *C. jejuni* PglF and S. enterica WbaP 4TMH domain. RMSD: 4.3 Å. Dehydratase domain omitted for clarity. **C.** Superposition of *C. jejuni* PglF and *S. enterica* WbaP DUF domain. RMSD: 2.3 Å. Dehydratase domain omitted for clarity.
